# Diverse fox circovirus (*Circovirus canine*) variants circulate at high prevalence in grey wolves (*Canis lupus*) from the Northwest Territories, Canada

**DOI:** 10.1101/2024.03.08.584028

**Authors:** Marta Canuti, Abigail V.L. King, Giovanni Franzo, H. Dean Cluff, Lars E. Larsen, Heather Fenton, Suzanne C. Dufour, Andrew S. Lang

**Affiliations:** Department of Veterinary and Animal Sciences, University of Copenhagen, Frederiksberg, 1870, Denmark; Department of Biology, Memorial University of Newfoundland, St. John’s, NL A1C 5S7, Canada; Department of Animal Medicine, Production and Health (MAPS), Padua University, 35020, Legnaro, Italy; Environment and Climate Change – North Slave Region, Government of the Northwest Territories, Yellowknife, NT X1A 2P9, Canada

## Abstract

Canine circoviruses (CanineCV) have a worldwide distribution in dogs and are occasionally detected in wild carnivorans, indicating their ability for cross-species transmission. However, fox circovirus, a lineage of CanineCV, has been identified exclusively in wild canids. We analyzed spleen samples from 159 grey wolves from the Northwest Territories, Canada, to investigate the molecular epidemiology of CanineCV and formulate hypotheses about virus ecology and evolution. Overall, 72 out of 159 (45.3%) animals tested positive. Virus prevalence was similar between males and females, adults and juveniles, and across the investigated years and locations. CanineCV infection was not associated with a poor body condition. While the percentage of co-infections with canine parvoviruses, investigated in a previous study, was high (63/72, 87.5%), the rate of parvovirus infection in CanineCV-negative animals was significantly lower (42/87, 48.3%, χ^2^ = 27.03, p < 0.001), and CanineCV infection was associated with a 7.5- and 2.4-fold increase in the risk of acquiring canine parvovirus 2 or canine bufavirus infections, respectively (odds ratios: 3.5-16.9 and 1.3-5.8). Although common risk factors cannot be ruled out, this suggests that CanineCV may facilitate parvoviral super-infections. Sequencing revealed high CanineCV genetic diversity, further exacerbated by recombination. Of the 69 sequenced strains, 87.5% were fox circoviruses, five were related to a fox circovirus-like recombinant strain, and one belonged to a distant lineage. In the phylogenetic analysis, the virus sequences were distributed according to sampling locations, with some viruses being geographically restricted. Different clades of viruses were identified in the same areas and over multiple years (2007-2019), indicating the co-existence of multiple endemic lineages in the investigated area. Phylogenetic analysis of all available complete fox circovirus genomes (32 from foxes and 15 from wolves from North America and Europe) demonstrated four lineages, each including sequences from this study. Within each lineage, strains segregated geographically and not by host. This implies that, although multiple lineages co-exist, viruses do not frequently move between locations. Finally, viruses from Europe and North America were mixed, indicating that the origin of the four lineages might predate the segregation of European and American wolf and fox populations. Given the high prevalence and diversity of fox circoviruses in wolves, these animals should be considered reservoir hosts for these viruses. Although we cannot exclude a lower susceptibility of dogs, the lack of fox circovirus in dogs could be due to environmental circumstances that prevented its spread to dogs. Given the high diversity and wild host specificity, we presume a long-lasting association between fox circovirus and canine hosts and hypothesize a higher likelihood of transmission from dogs to wild animals than vice versa. Further studies should investigate other sympatric wild species and additional locations to explore the possible existence of additional maintenance hosts and the reasons behind the marked difference in cross-species transmission dynamics among CanineCV lineages.

## Introduction

Circoviruses (family *Circoviridae*, genus *Circovirus*) are small viruses with circular, covalently closed, single-stranded DNA genomes of approximately 2 Kb. The circoviral genome is ambisense and contains two major open reading frames (ORFs), coding for the replication-associated (Rep) and the capsid (Cap) proteins. The 5′ ends of the two ORFs are separated by a non-coding region including a stem-loop structure with a nonanucleotide motif marking the origin of DNA replication (Breitbart et al., 2017).

Circoviruses have been identified in many terrestrial and aquatic mammals, including most families of the order Carnivora, such as ursids (Alex et al., 2020), viverrids (Nishizawa et al., 2018), mustelids (Lian et al., 2014; Bandoo et al., 2021), herpestids (Gainor et al., 2021), felids (Payne et al., 2020; Cerna et al., 2023), pinnipeds (Chiappetta et al., 2017; Patterson et al., 2021), and canids (Kapoor et al., 2012; Bexton et al., 2015). Although 60 circoviral species have currently been defined, the pathogenic role and ecology of these viruses, including their potential host ranges, have been only rarely investigated as most of them have been discovered within the framework of metagenomic investigations.

The canine circovirus (CanineCV, species *Circovirus canine*) is the sole member of the genus containing viruses currently known to infect canines. CanineCV was first identified in 2012 in serum and tissue samples of North American dogs (Kapoor et al., 2012; Li et al., 2013), and subsequent epidemiological investigations have confirmed its global presence in dogs (Gomez-Betancur et al., 2023). In these studies, clinical signs indicative of gastroenteritis, respiratory disease, and hematological and immune disorders have been found associated with CanineCV infection in domestic dogs. However, no conclusive evidence has yet proven the involvement of this virus, alone or as a co-infecting agent, in clinical disease (Gomez-Betancur et al., 2023). Nonetheless, some circoviruses, such as porcine circovirus 2 (PCV-2) or beak and feather disease virus (BFDV), are known animal pathogens and are of high veterinary importance, especially considering their capacity to induce marked immunosuppression (Fehér et al.). These viruses are also considered a threat to wildlife (Kasimov et al., 2023). Therefore, the clinical importance of CanineCV should not be underestimated.

The same strains of CanineCV that circulate in domestic animals have also been identified in wild carnivorans, such as wolves, badgers, foxes, and jackals (Zaccaria et al., 2016; Arcangeli et al., 2020; Balboni et al., 2021; de Villiers et al., 2023), indicating the occurrence of cross-species transmission among sympatric wild species that is presumably also enabled by contacts between domestic and wild carnivorans. A particular clade of viruses within the species *Circovirus canine*, known as fox circovirus, is notable because it includes genetically distant viruses that were considered a separate species in the past. However, it is now recognized that fox circoviruses do not fulfill the genetic requirements to obtain a distinct taxonomic status and remain part of *Circovirus canine* (Breitbart et al., 2017). Nonetheless, it is intriguing that fox circoviruses have only been identified in wild canids (foxes and wolves) across Europe and North America and, as of now, never in dogs. In particular, the first fox circovirus was detected in red foxes with meningoencephalitis in the United Kingdom (Bexton et al., 2015) and later on, fox circoviruses were also identified in Croatian red foxes (Lojkić et al., 2016), in red foxes and wolves from Italy (Balboni et al., 2021; Franzo et al., 2021), in Arctic foxes from Svalbard (Urbani et al., 2021), in red foxes from Northern Norway (Urbani et al., 2021), and in red and Arctic foxes from Newfoundland and Labrador, Canada (Canuti, Rodrigues, et al., 2022).

Despite their uncertain clinical significance, given their peculiar host distribution pattern, these viruses offer an interesting perspective to study circovirus transmission dynamics and help understand viral transmission dynamics between wild and domestic animals. In this study, we explored samples collected from Canadian grey wolves (Northwest Territories) (Canuti, Fry, et al., 2022) to investigate the molecular epidemiology of CanineCV in this previously unexplored region. Additionally, through comprehensive full genome sequence analyses, the study also aimed to explore the international spread of fox circovirus and formulate hypotheses about its ecological patterns and evolutionary pathways.

## Material and methods

### Samples

This study included samples of DNA previously isolated from the spleens of 159 gray wolves from various locations within the Northwest Territories, Canada (Canuti, Fry, et al., 2022) (Figure 1). Wolf carcasses were previously processed as part of ongoing scientific monitoring procedures by the Department of Environment and Natural Resources (now Environment and Climate Change) of the Government of the Northwest Territories (GNWT) under research permits WL003091, WL005627, WL005634, WL005622, WL005761, WL005768, WL500098, WL500199, WL500289, WL500393, WL500481, WL500666. Handling of samples at Memorial University of Newfoundland was done under Animal Use Protocol 20-04-AL.

**Figure 1.**
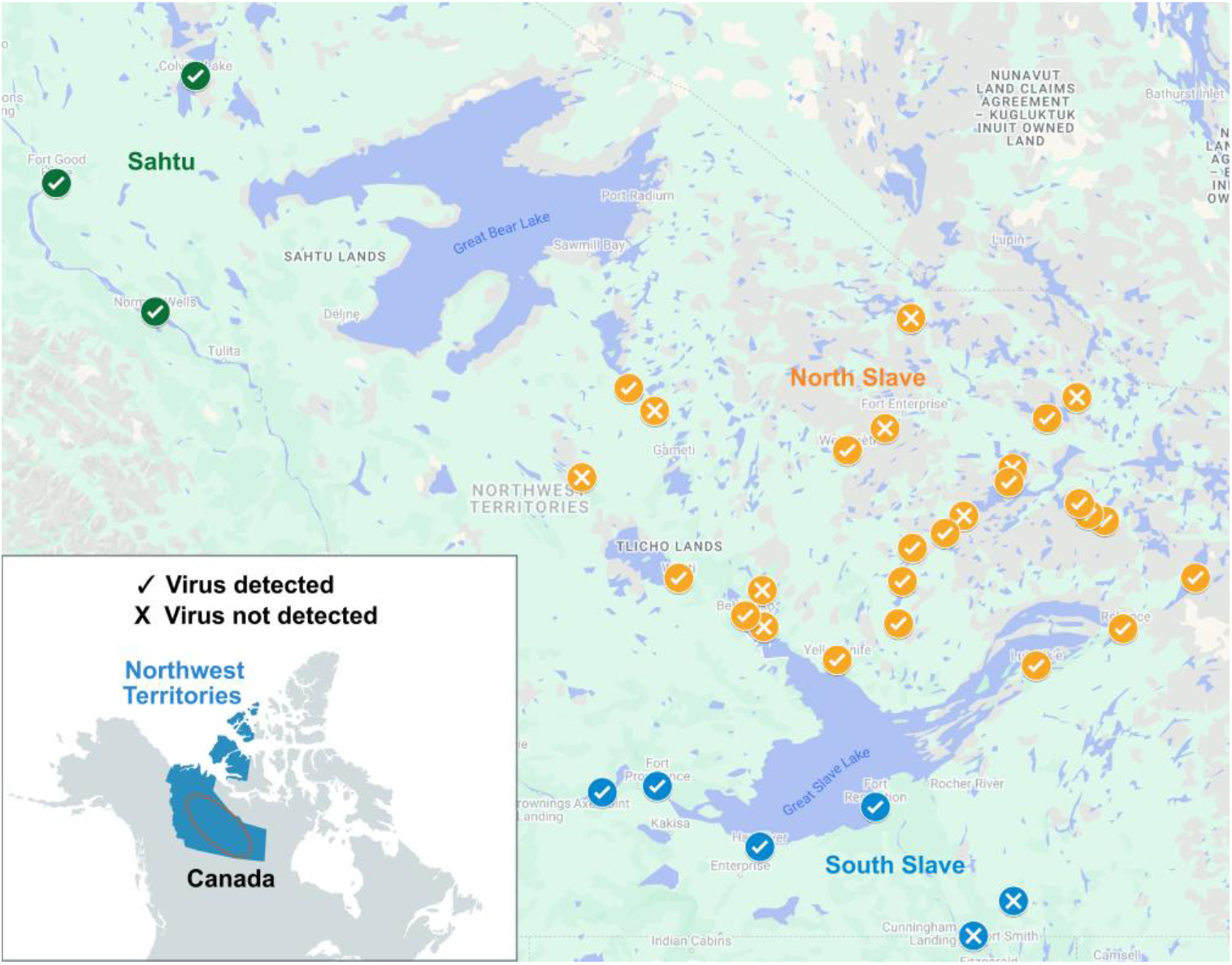
Sample collection locations. The map on the bottom-left shows the location of the Northwest Territories within Canada. The main map shows an enlargement of the circled region. Approximate locations of sample collection are indicated by circles (each location could include multiple samples) depicting a ✓ in case of virus detection and an X in case of lack of virus detection. The three different investigated regions have been color-coded: green for Sahtu, orange for the North Slave Region, or blue for the South Slave Region. The maps were created with MapChart © and Google Map (Map data ©2019 Google)

Harvest locations were recorded for all samples and three main sampling areas were defined using large lakes as separation landmarks: west of Great Bear Lake (Sahtu, N = 7), the zone between Great Slave Lake and Great Bear Lake (North Slave Region, NSR, N = 140), and the area south of Great Slave Lake (South Slave Region, SSR, N = 12) (Figure 1). Samples were collected between 2007 and 2019 (2007: N = 17, 2008: N = 19, 2009: N = 5, 2010: N = 24, 2011: N = 16, 2012: N = 2; 2013: N = 5, 2014: N = 1, 2016: N = 6, 2017: N = 1; 2018: N = 3, 2019: N = 60), predominantly in winter (154/159: 96.9%). Sixty-five animals were female (48.9%) and 94 were male (59.1%), while the age was recorded only for 70 animals, including 8 juveniles, defined as animals ≤2 years of age, and 62 adults. Body conditions were scored for 64 animals based on internal and external body condition ranks (score of 0-4 with ≥2 considered adequate). Finally, the positivity of these animals to five species of dog parvoviruses (canine parvovirus 2: CPV-2, canine bufavirus: CBuV, cachavirus: CachaV, minute virus of canines: MVC, canine bocavirus 2: CBoV-2) was known from a previous investigation. Animal information details are described in (Canuti, Fry, et al., 2022).

### Screening and sequencing

Specific primers designed to detect all currently known viruses within the species *Circovirus canine* were designed for this study and used for screening DNA samples through a hemi-nested PCR. For the first PCR (35 cycles), 2.5 μl of DNA were used as input, and primers CCV_F1 (ARCGNAATGACTTACATGATGC) and CCV_R1 (GCGAGAGGCCTTTATYTYTC) were used to amplify a 578-nt portion of the Rep ORF; primers CCV_F2 (TGGTGGGAYGGYTACGATGG) and CCV_R1 were used during the nested step (25 cycles) to amplify a fragment of 324 nt. The complete genomic sequences of the strains with the highest viral loads (those that were already positive after the first PCR) were obtained through sequencing two overlapping PCR products obtained with primer pairs CCV_F4a (CTTGTRGCATTCCCTCTTCC) / CCV_R1 (1475 nt) and CCV_F7 (TTGGTAACAAGGAYGCCCTC) / CCV_R7 (CAGCATCTCCCACCRTTCAG) (803 nt). The addition of DMSO (5.2%) was required for the amplification of the fragment included between primers CCV_F7 and CCV_R7. All PCRs were run in a 25 μl reaction volume at an annealing temperature of 50°C and using the DreamTaq Green PCR Master Mix (Thermo Fisher Scientific). All obtained amplicons were purified with AMPure XP beads (Agencourt) and outsourced for Sanger sequencing.

### Sequence analyses

Assembly of obtained sequences and open reading frame (ORF) predictions were performed in Geneious R11 (Biomatters). Reference sequences available in the GenBank database (accessed in December 2023) were downloaded and used for comparisons (accession numbers available in Supplementary Figure S1 in the Appendix). Sequence alignments were performed with MAFFT (Katoh & Standley, 2013), and maximum-likelihood phylogenetic trees were inferred with IQ-TREE 2 (Minh et al., 2020) using the substitution model with the lowest Bayesian information criterion calculated by ModelFinder (Kalyaanamoorthy et al., 2017). Branch support was determined with ultrafast bootstrapping (Hoang et al., 2018) and SH-like approximate likelihood ratio test (SH-aLRT) (Guindon et al., 2010). Recombination analyses were performed with RDP version 5.53 (Martin et al., 2021) and SimPlot (Salminen et al., 1995).

### Statistical analyses

Categorical variables were expressed in percentages with relative 95% normal intervals (95% intervals of confidence, 95% IC). Proportions were compared using the Chi-square (χ^2^) test or Fisher’s exact test when appropriate. Two-sided p-values < 0.05 were considered statistically significant. Simple and multiple logistic regression models were fitted to evaluate CanineCV positivity as a risk factor for poor body conditions (internal body score or an average of internal and external body score <2, with CPV-2 and CBuV included as co-variates) and for parvovirus infection, both overall and for single parvoviruses (with all other parvoviruses included are co-variates). Odds ratios (OR) with relative 95% IC were considered as the measure of effect and precision, respectively, while the Wald test was used to assess the significance of the regression beta coefficient. Analyses were performed with Past version 4.08 (Hammer et al., 2001) and JASP 0.17.1 (JASP Team, 2023).

## Results

### CanineCV epidemiology among grey wolves

Out of the 159 samples, 72 tested positive (prevalence: 45.3%, 95% CI: 37.6-53.0%). The prevalences were similar between males and females (44/94, 46.8% vs. 28/65, 43.1%; χ^2^ = 0.21, p = 0.6), adults and juveniles (22/62, 35.5%, vs. 5/8, 62.5%; Fisher’s exact: p = 0.3), and among the investigated years (27/65, 41.5% in 2007-2010, 14/24, 62.5% in 2011-2015, and 31/70, 44.3% in 2016-2019; χ^2^ = 2.05, p = 0.4), months (13/22, 59.1% in January, 15/37, 40.5% in February, 27/59, 45.8% in March, 6/9, 66.7% in November, and 9/27, 33.3% in December; χ^2^ = 5.25, p = 0.3), and locations (63/140, 45.0% in NSR, 5/7, 71.45% in Sahtu, and 4/12, 33.3% in SSR, Fisher’s exact: p = 0.3). Interestingly, the viral prevalence was higher around the capital city of Yellowknife (NSR, 13/16, 81.3%). Finally, CanineCV positivity was not identified as a risk factor for poor internal body conditions (non-adjusted OR: 0.9 (0.3-3.3); adjusted OR: 1.1 0.3-4.7)) and average internal and external poor body conditions (non-adjusted OR: 0.4 (0.1-1.2); adjusted OR: 0.4 (0.2-1.1)).

These animals were previously investigated for the presence of five species of dog parvoviruses (Canuti, Fry, et al., 2022). A comprehensive list of the prevalence of the detected viruses as well as the number of co-determinations for each virus in the investigated population is shown in Figure 2. Interestingly, as shown in Table 1, the percentage of parvovirus-positive animals was significantly higher in the group of CanineCV-positive animals (63/72, 87.5%) compared to the CanineCV-negative animals (42/87, 48.3%, χ^2^ = 27.025, p < 0.001), potentially indicating a synergistic effect between CanineCV and parvovirus infection. This was also true when evaluating the association between CanineCV and the two parvoviruses with the highest prevalence, CPV-2 (χ^2^ = 32.49, p < 0.001) and CBuV (χ^2^ = 6.96, p = 0.008). Finally, CanineCV infection was associated with a more than seven-fold increase in the risk of acquiring CPV-2 infection (non-adjusted OR: 7.5 (3.6-15.7); adjusted OR: 7.7 (3.5-16.9)) and with a more than two-fold increase in acquiring CBuV infection (non-adjusted OR: 2.4 (1.2-4.5); adjusted OR: 2.8 (1.3-5.8)). No correlation was found between CPV-2 and CBuV infections.

**Table 1.** Evaluation of CanineCV infection as a risk factor for acquiring a parvovirus super-infection.

|  | CanineCV + (N=72) |  | CanineCV - (N=87) |  |  | Odds ratio<sup>a</sup> |  |  |
| --- | --- | --- | --- | --- | --- | --- | --- | --- |
|  | N | % | N | % | P | OR | 95% IC | P |
| CPV-2 | 44 | 61.1 | 15 | 17.2 | < 0.001 | 7.5 | 3.6-15.7 | < 0.001 |
| CBuV | 39 | 54.2 | 29 | 33.3 | 0.008 | 2.4 | 1.2-4.5 | 0.009 |
| CachaV | 4 | 5.6 | 1 | 1.2 | 0.1 | 5.1 | 0.6-46.3 | 0.2 |
| CBoV-2 | 5 | 6.9 | 3 | 3.5 | 0.5 | 2.1 | 0.5-9.1 | 0.3 |
| MVC | 1 | 1.4 | 0 | 0.0 | 0.5 | / | / | / |
| Any parvovirus | 63 | 87.5 | 42 | 48.3 | < 0.001 | 7.5 | 3.3-17.0 | < 0.001 |

Note: CanineCV: canine circovirus; CPV-2: canine parvovirus 2; CBuV: canine bufavirus; CachaV: cachavirus; MVC: minute virus of canines; CBoV-2: canine bocavirus 2; N: number of positive samples; %: prevalence, OR: odds ratio for CanineCV infection; IC: interval of confidence on OR; /: number of positive samples too low

**Figure 2.**
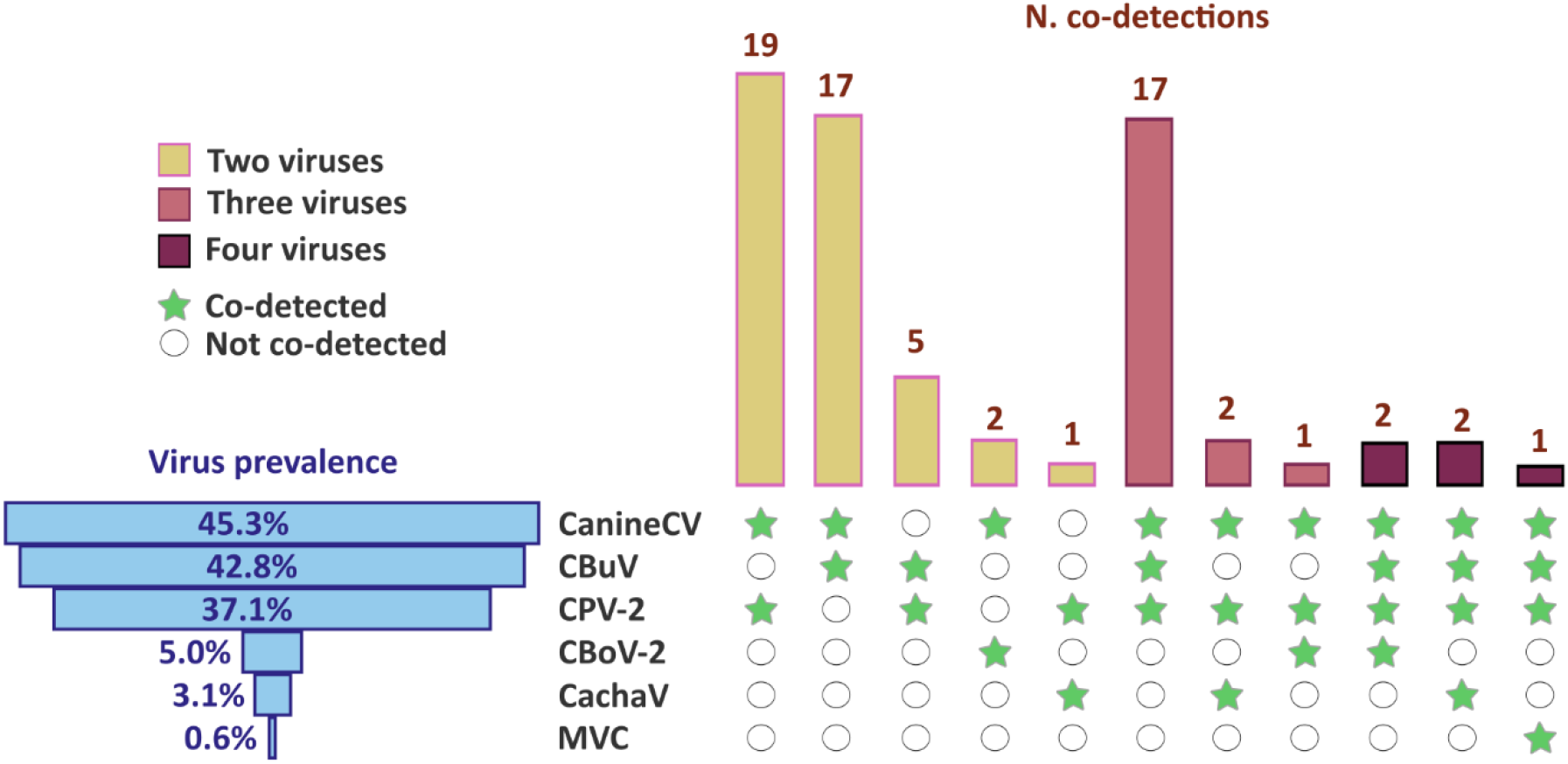
Identified co-detections in the investigated population. The bars on the lower left represent the overall prevalence of each virus, while the bars on the top right represent the number of co-detections involving the viruses indicated by green stars. Co-detections are color-coded according to the number of identified viruses, as explained in the legend on the left. CanineCV: canine circovirus; CPV-2: canine parvovirus 2; CBuV: canine bufavirus; CachaV: cachavirus; MVC: minute virus of canines; CBoV-2: canine bocavirus 2

### CanineCV diversity among grey wolves

The amplicons obtained through the screening PCRs were all subjected to Sanger sequencing to molecularly type the viruses. Sequences obtained from three of the positive animals clearly showed the presence of multiple circoviral variants, as demonstrated by the presence of several polymorphic sites (co-occurring peaks), while 69 viruses could be successfully typed. As shown in Figure 3, six animals were infected with variants within highly supported clades containing viruses also detected in dogs. Five of these (clade A) clustered together with a virus identified once in a dog from California (UCD3-478) (Li et al., 2013) and thereafter recognized as a recombinant between a fox circovirus and a virus from another clade (Canuti, Rodrigues, et al., 2022). Nonetheless, the 5 viruses from this study within this clade formed a highly-supported monophyletic sub-clade. The other one (clade G) was part of a clade comprising several viruses identified in dogs worldwide (China, Viet Nam, Iran, Italy, Germany, Namibia, USA, Argentina, Colombia, Brazil) as well as in wildlife (wolf, badger and fox in Italy, and jackal in Namibia) (Supplementary Figure S1 in the Appendix, support: 97.2/98). All other viruses (87.5%) could be classified with high support as fox circovirus. These were distributed into five different clades (B-F), with clades E and F comprising the majority (72.5%) of the identified strains.

**Figure 3.**
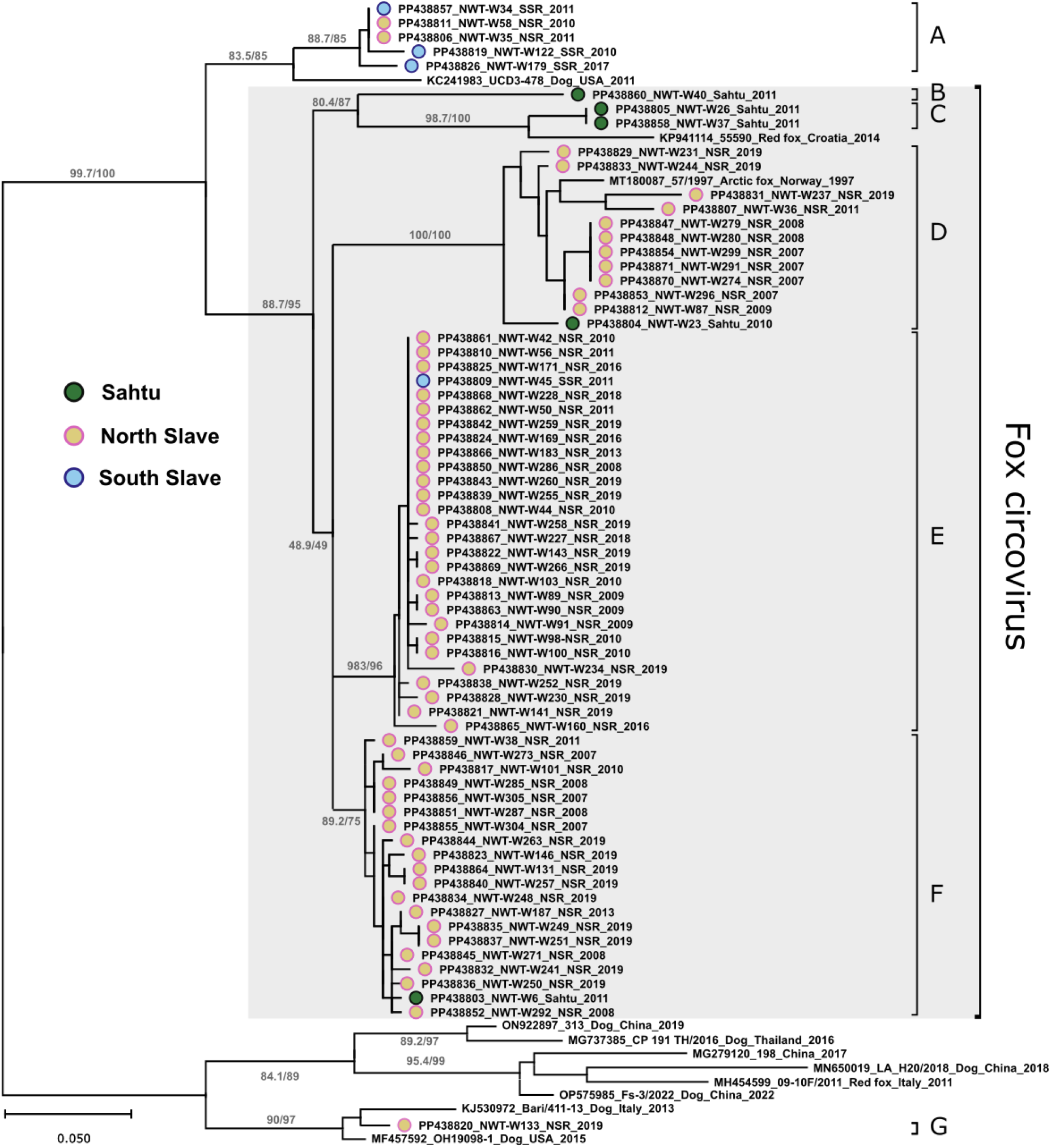
Molecular epidemiology of fox circovirus in wolves of the Northwest Territories, Canada. The tree shows the phylogenetic relationships between sequences of canine circoviruses from this study and some reference sequences representing each major clade of the species *Circovirus canine*. The tree, based on a 265-nt alignment of the Rep gene, was built with the maximum-likelihood method based on the TN+F+G4 model with IQ-Tree. The outcomes of the SH-aLRT and bootstrap tests (1000 replicates) are shown for the main nodes and branch lengths are proportional to genetic distances as indicated by the scale bar. Strains identified in this study are labeled with a dot colored based on the sampling location (blue: South Slave Region (SSR); orange: North Slave Region (NSR); green: Sahtu). Following the GenBank accession numbers, sequences are indicated by the strain name followed by the host (for reference sequences only), sampling location, and collection year. The tree built with all reference sequences is available in Supplementary Figure S1 in the Appendix.

Although a smaller number of samples was available from two of the three investigated areas (Figure 1), a certain clustering based on sampling location could be observed (Figure 3). Three out of four viruses from SSR were included in clade A, indicating a predominance of viruses from this clade in that location. Three out of five viruses from Sahtu were in clades B and C, which included only viruses from this region. Another Sahtu virus clustered within another clade (D) that otherwise exclusively contained viruses from NSR, although in a separate and basal branch. Finally, viruses from NSR were distributed across five clades (i.e., A, D, E, F, and G), none of which was specific to this area. All three co-infected animals were from NSR. While clades A, D, E, and F included multiple viruses sampled over various years, suggestive of endemicity, the strain in clade G was unique and possibly not associated with onward transmission. No patterns that related the year of sampling with tree topology were observed.

### Global epidemiology of fox circoviruses

A total of 15 complete genomic sequences from circoviruses identified in this study were obtained. These represented all the fox circovirus and recombinant clades identified in this study. Unfortunately, we could not amplify the genome of the virus from clade G because of the low viral load.

Obtained sequences were compared to 271 sequences from members of the species *Circovirus canine* available in public repositories. As expected, in the maximum likelihood phylogenetic tree all these sequences clustered within a highly supported clade (red branches in Figure 4) including the sequences UCD 478 and NWTW-34 (the recombinant strains) in a basal branch and all 47 so far completely sequenced fox circovirus sequences grouped together. Complete genomes were 89.0-99.9% identical to each other and showed a consistent genome organization. Rep proteins were more conserved (89.4-100%) compared to Cap proteins (87.4-100%). Interestingly, the final 23 amino acids of the Rep protein had a higher variability compared to the rest of the protein.

**Figure 4.**
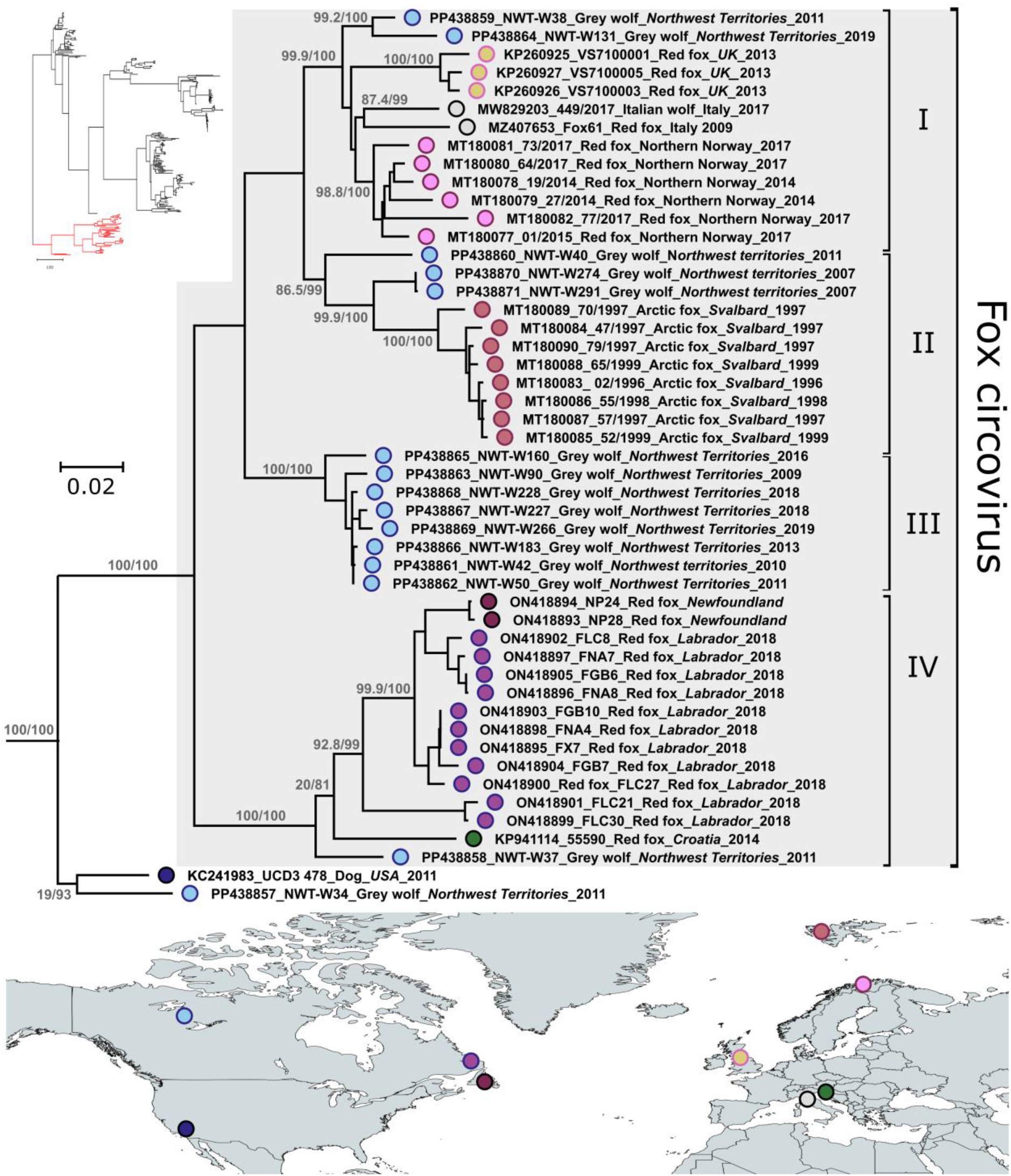
Global molecular epidemiology of fox circovirus. The tree shows the phylogenetic relationships among fox circoviruses identified in various parts of the world. The tree is based on complete genome sequences and was built with the maximum-likelihood method with the TIM3+F+I+G4 model with IQ-Tree. The outcomes of the SH-aLRT and bootstrap test (1000 replicates) are shown for the main nodes and branch lengths are proportional to genetic distances as indicated by the scale bar. The tree on the top-left represents the global *Circovirus canine* phylogeny and red branches indicate the clade that is shown magnified on the right. Strains are labeled based on the collection site as shown in the map below (light blue: Northwest Territories; dark blue: California; purple: Labrador; magenta: Newfoundland; orange: UK; dark pink: Svalbard; light pink: mainland Norway; grey: Italy; green: Croatia). Following the GenBank accession numbers, sequences are indicated by the strain name followed by host, sampling location, and collection year. The map was created with MapChart ©

Fox circoviruses could be divided into four highly supported major clades, indicated by Roman numerals in Figure 4 and Supplementary Table S1 in the Appendix. Interestingly, sequence NWT-W40 was identified as a likely recombinant virus, containing a small (nt 780-1060) insert with low identity to any identified strain, included within a backbone that was mostly clade II-like (Supplementary Figure S2 in the Appendix). This also explains the different clustering of this sequence in the two phylogenetic trees shown in Figures 3 and 4. Additionally, sequence NWT-W34 was again identified as recombinant, together with UCD 478, as previously reported (Canuti, Rodrigues, et al., 2022).

Because of its intriguing epidemiology, the global distribution and phylogeny of fox circovirus sequences were examined in detail. All fox circoviruses identified so far were detected in wildlife, with 32 strains being from foxes (red foxes (*Vulpes vulpes*) from Newfoundland and Labrador in Canada, Croatia, Italy, United Kingdom, and Arctic fox (*Vulpes lagopus*) Svalbard) and 15 from wolves (Italian wolf (*Canis lupus italicus*) in Italy and grey wolf (*Canis lupus*) from the Northwest Territories). Sequences from the Northwest Territories were the only ones represented in each of the identified clades. Viruses from Norway showed the highest diversity after those from this study, being identified in two different clades, although one clade was unique to one of the islands of Svalbard and the other one was unique to mainland Norway. Although there was only one location where viruses from both wolves and foxes were present, we could observe that the strains clearly segregated geographically and not by host, with viruses from the same location consistently forming monophyletic subclades. In fact, the same host species were infected with strains belonging to different clades (e.g., strains from clades I, II, and III detected in grey wolf), and viruses of the same clade infected multiple hosts (e.g., clades II and III including viruses from wolf and fox).

## Discussion

Despite its relatively recent discovery (Kapoor et al., 2012; Li et al., 2013), several studies have investigated the molecular epidemiology of CanineCV in dogs, demonstrating its high diversity and worldwide distribution (Gomez-Betancur et al., 2023). On the contrary, only a few studies have evaluated the presence of this virus in wild animals. Studying viruses in wild animals does not only have relevance for wildlife conservation, but it is also important in terms of domestic animal health as wild animals can be reservoir hosts for pathogens that can spill over into new hosts and negatively impact their health (Martin et al., 2011). CanineCV strains have so far been identified in a few wild carnivorans (wolves, badgers, foxes, and jackals) and the phylogeny of the various strains is driven by their geographic origin, rather than the host source. This is an indication that cross-species transmission has occurred several times, in many different locations (Zaccaria et al., 2016; Arcangeli et al., 2020; Urbani et al., 2021; Balboni et al., 2021; Franzo et al., 2021; Canuti, Rodrigues, et al., 2022; de Villiers et al., 2023). Nonetheless, endemicity and persistent transmission chains within wild animal populations have also been documented (de Villiers et al., 2023).

### CanineCV is endemic among wolves of Northern Canada

In our study, 45% of the investigated animals were CanineCV-positive, indicating endemic circoviral circulation among wolves of the Northwest Territories. This number is in line with previous studies that also identified high infection prevalence in wild animals (Bexton et al., 2015; Canuti, Rodrigues, et al., 2022; de Villiers et al., 2023). Prevalence estimates could be affected by the type of analyzed sample, explaining potential variation across studies (Canuti, Rodrigues, et al., 2022). In this study, we used DNA isolated from spleens. While the presence of a virus in the spleen indicates a systemic infection, it does not necessarily reflect an acute infection stage, as it can be the outcome of viral or DNA persistence. Indeed, the most studied circovirus, PCV-2, is known to cause chronic infections, with the virus persisting for months after the initial infection and being detectable in blood for long periods of time (Shibata et al., 2003; Opriessnig et al., 2010). Nonetheless, our previous investigation showed that, in foxes, positivity rates for CanineCV did not significantly differ between stool and spleen samples (Canuti, Rodrigues, et al., 2022).

Also consistently with our previous findings in Canadian foxes (Canuti, Rodrigues, et al., 2022), we identified a high circoviral diversity among the investigated animals and the occurrence of recombination. However, differently from foxes, viruses identified in wolves were not monophyletic and belonged to three different CanineCV lineages and seven clades. Of these, viruses from four different clades were detected in different areas and spanning multiple years (2007-2019), indicating the co-existence of multiple endemic viral lineages across large distances. Some strains were unique to Sahtu, which is distant from the other two regions (SSR and NSR) that are closer together and more interconnected. Interestingly, however, strains from SSR were mostly included in only one of the clusters. This pattern may be the consequence of bottleneck and geographical isolation and/or behavioral separation, resulting in some strains being geographically restricted.

Finally, for one virus we found no evidence for spreading, although this was identified in a sample collected in the final year of the study. This virus was part of a cluster that, unlike others, had been previously detected in both dogs and wild animals, and we could speculate that this was a case of spillover from a different host species with limited or no onward circulation. For this reason, investigating the origin of this specific virus would be interesting. Regrettably, samples from sympatric species were unavailable for this purpose.

### Does CanineCV facilitate a secondary parvoviral infection?

One of the most interesting findings of our study was the intriguing relationship between CanineCV and highly prevalent parvoviruses, which were investigated in the same wolf population in a different study (Canuti, Fry, et al., 2022). A parvoviral co-infection was detected in 87.5% of CanineCV-positive animals, and these mostly involved the two parvoviruses with the highest prevalences, CBuv and CPV-2.

Remarkably, the percentage of CPV-2- and CBuV-positive animals was significantly higher (p < 0.001) among CanineCV-positive animals compared to negative ones (48.3%). Additionally, CanineCV infection was associated with more than seven-fold increase in the risk of acquiring a CPV-2 infection (p < 0.001) and with more than two-fold increase in the risk of acquiring a CBuV infection (p = 0.009). These results could indicate a possible synergistic effect between CanineCV and parvoviral infections, where CanineCV infection favors a parvoviral co-infection, that co-infection with parvovirus triggers the CanineCV virus to persist, or *vice versa*. However, the presence of common infection risk factors cannot be excluded either. While CPV-2 is known to cause immune suppression and alterations of the enteric mucosa that could favor super-infections (Shahbazi Asil et al., 2023), the effects of CBuV and CanineCV on the immune system are unknown. However, some well-studied circoviruses, such as PCV-2, can induce marked immunosuppression and favor the replication of other pathogens (Fehér et al.), and co-infections of PCV-2 with porcine parvovirus (PPV) are also frequently detected (Kim et al., 2003).

Given the data available from this study, it is not possible to establish which one was the initial infection; however, it was noticeable that no correlation was found between CPV-2 and CBuV-positive animals. This may indicate that an earlier CanineCV infection leads to a predisposition to super-infections by other pathogens, maybe also mediated by immunosuppression, as it has also been hypothesized in other studies (Dankaona et al., 2022). Interestingly, a recent investigation showed that CanineCV infection increases the replication level of CPV-2 *in vitro* (Hao et al., 2022) and our epidemiological data might support those findings and extend them to CBuV. Undoubtedly, further investigations are required to confirm this hypothesis. However, our analysis revealed no significant link between CanineCV infection and poor body condition, a finding that is also consistent with what was identified for CPV-2 in the same population (Canuti, Fry, et al., 2022). The apparent increased resilience to symptomatic clinical courses in these wild animals warrants further investigation through dedicated studies.

### Viral hosts and ecology

Not all CanineCV clades are alike, and strains classified as “fox circovirus” demonstrate a peculiar epidemiological profile. Unlike all other CanineCV clades, fox circoviruses have so far been identified exclusively in wild animals, specifically foxes and wolves (Franzo et al., 2021; Canuti, Rodrigues, et al., 2022). In our population, out of the 69 successfully typed strains, 87.5% belonged to the fox circovirus group, resulting in a prevalence specific for fox circovirus of 37.7%. An additional five strains belonged to a clade of viruses that likely originated after recombination between a fox circovirus and a CanineCV from another clade (Canuti, Rodrigues, et al., 2022). A virus in this clade was also identified once in a dog (Li et al., 2013). Despite the unavailability of samples from sympatric species and the possible existence of additional maintenance wild hosts preventing us from making assumptions generalizable to the whole region, fox circoviruses and highly genetically related viruses appear the most common CanineCV strains in wild wolves from this region of Northern Canada. Indeed, only one virus was identified in a different part of the phylogenetic tree.

It is currently unclear whether the tight association between fox circoviruses and wildlife is a consequence of a particular viral phenotypic feature and host adaptation or whether it is simply related to epidemiological factors preventing or limiting effective contacts with domestic dogs (Franzo et al., 2021; Canuti, Rodrigues, et al., 2022). However, viral endemicity in wolves, which belong to the same species as domestic dogs, suggests that fox circoviruses are likely capable of infecting dogs. The lack of identification in dogs is thus likely either the result of under-sampling or of environmental factors.

Our current findings, along with data from existing literature, suggest that the likelihood of CanineCV transmission from domestic dogs to wild animals is greater than the reverse scenario of transmission from wild animals to dogs. Indeed the likelihood of cross-species transmission between sympatric species may differ for different viruses and hosts (Canuti et al., 2020). However, it is important to note that no samples from dogs were available for our investigation and that viral diversity in wildlife is largely understudied, so these conclusions could be biased by sample and sequence availability.

While fox circoviruses have so far only been identified in foxes and wolves and only in Europe and North America, they may also have a broader host and geographic distribution. Overall, four different fox circovirus lineages have been identified and only two locations were positive for viruses from more than one lineage. Specifically, two lineages were found in Norway, although one lineage was unique to continental Norway and another one to an island of the Svalbard archipelago (Urbani et al., 2021), and viruses from all four lineages were identified in the Northwest Territories. Within each lineage, viruses clustered geographically indicating that, although multiple lineages can co-exist in one location, viruses tend to differentiate locally and do not (frequently) move between different locations. Being restricted to wild animals, these viruses may not benefit from the remarkable mobility of domestic dogs, as happens for other canine viruses (Mira et al., 2019; Alfano et al., 2022; Franzo et al., 2023), explaining the more constrained distribution and stronger geographical clustering. Nonetheless, when comparing the data we obtained here to those from our previous study performed with foxes from Labrador (Canuti, Rodrigues, et al., 2022), it is noteworthy that the viruses identified in these two regions differ significantly, highlighting a higher viral diversity in wolves from the Northwest Territories. Finally, strains from very distant locations were intermingled together in the tree, indicating that the origin of the four fox circovirus clades possibly pre-dates the segregation of European and American wolf and fox populations. However, the same pattern could also be explained by viral transfer by yet undetected routes and further investigations examining additional hosts and locations are required to verify this hypothesis.

## Conclusions

Given the high prevalence and the high diversity of fox circoviruses in wolves, their role as reservoir hosts for this virus can be stated, and a long-lasting virus-host association can be hypothesized. However, future studies may evidence the existence of additional maintenance hosts. While the clinical impact of these viruses on wild and domestic animals is debatable, circoviruses may enhance or facilitate the infection of other viruses, particularly CPV-2 and CBuV, with relevant implications for wild animal health. Intriguingly, fox circovirus, which is widespread across wild animal populations, has never been detected in dogs and further studies should assess the presence of this virus among dogs and other wild sympatric animal populations to clarify its ecology and transmission dynamics. The marked difference in cross-species transmission dynamics between viruses belonging to different CanineCV clades is also investigation-worthy. Given the high diversity and geographic segregation of fox circoviral strains, it is conceivable that this virus has been circulating among wild carnivorans for a long time and it is likely that future research involving additional carnivoran species and locations will reveal an even greater genetic diversity and wider host range.

## Appendix

**Supplementary Table S1.** Between and within (Italics) clade pairwise sequence identities, expressed as a percentage of 1-p distance. Clades correspond to those shown in Figure 4.

|  | Clade I | Clade II | Clade III | Clade IV |
| --- | --- | --- | --- | --- |
| **Clade I** | *94.1-99.3* |  |  |  |
| **Clade II** | 88.7-91.6 | *93.4-99.9* |  |  |
| **Clade III** | 91.4-93.8 | 90.1-91.4 | *97.6-99.9* |  |
| **Clade IV** | 89.1-91.9 | 91.9-94.6 | 91.6-93.1 | *93.1-99.9* |

**Supplementary Figure S1.**
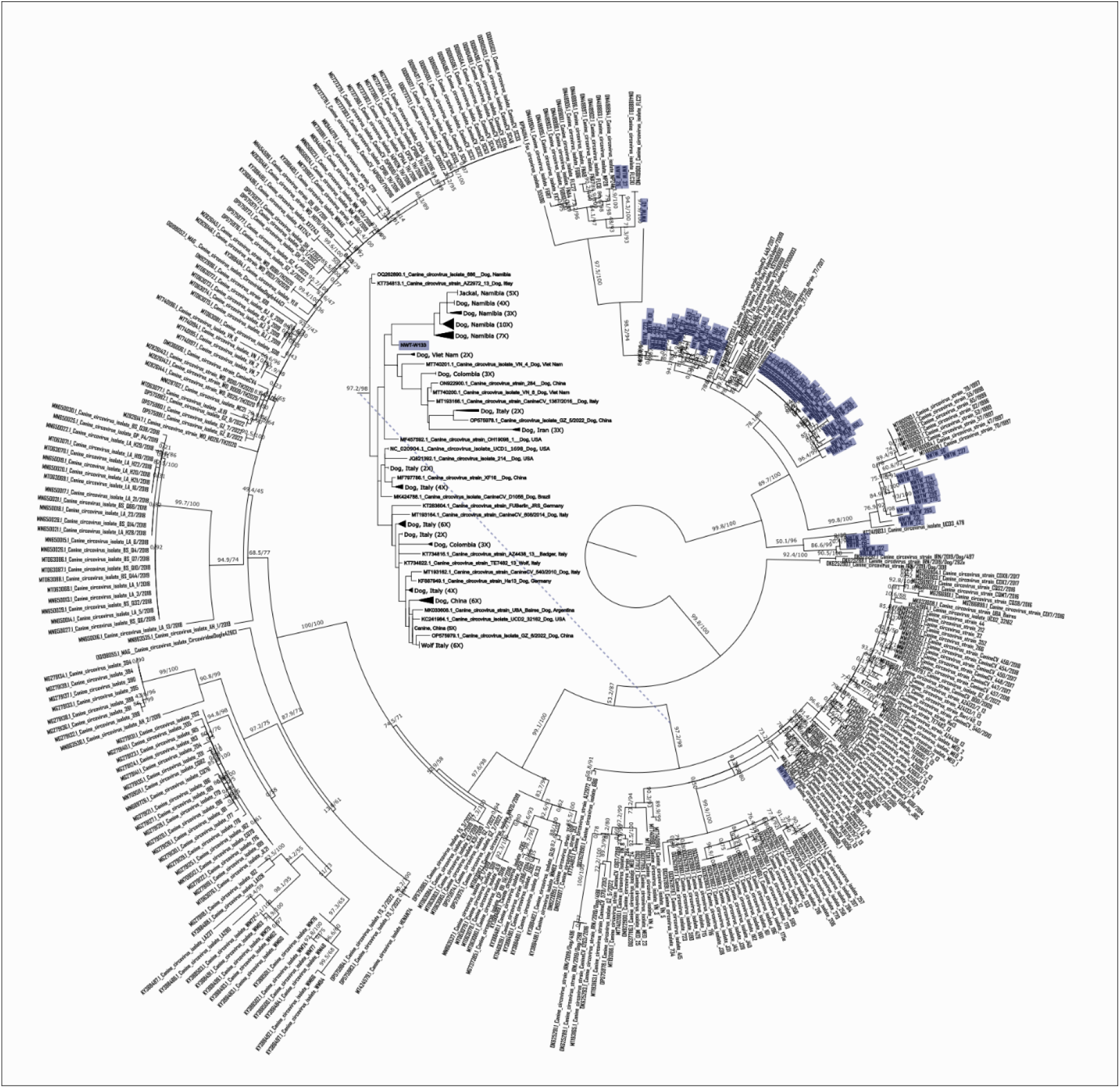
Relationships between canine circoviruses of wolves of the Northwest Territories, Canada, with other *Circovirus canine* strains reported in the literature. The tree, based on a 265-nt alignment of the Rep gene, was built with the maximum-likelihood method based on the TIM3+F+I+G4 model with IQ-Tree. The outcomes of the SH-aLRT and bootstrap tests (1000 replicates) are shown for all nodes and branch lengths are proportional to genetic distances. The strains identified in this study are highlighted in purple. The branch including strain NWTW-133 is enlarged in the middle, where information about the host and location of reference strains is indicated.

**Supplementary Figure S2.**
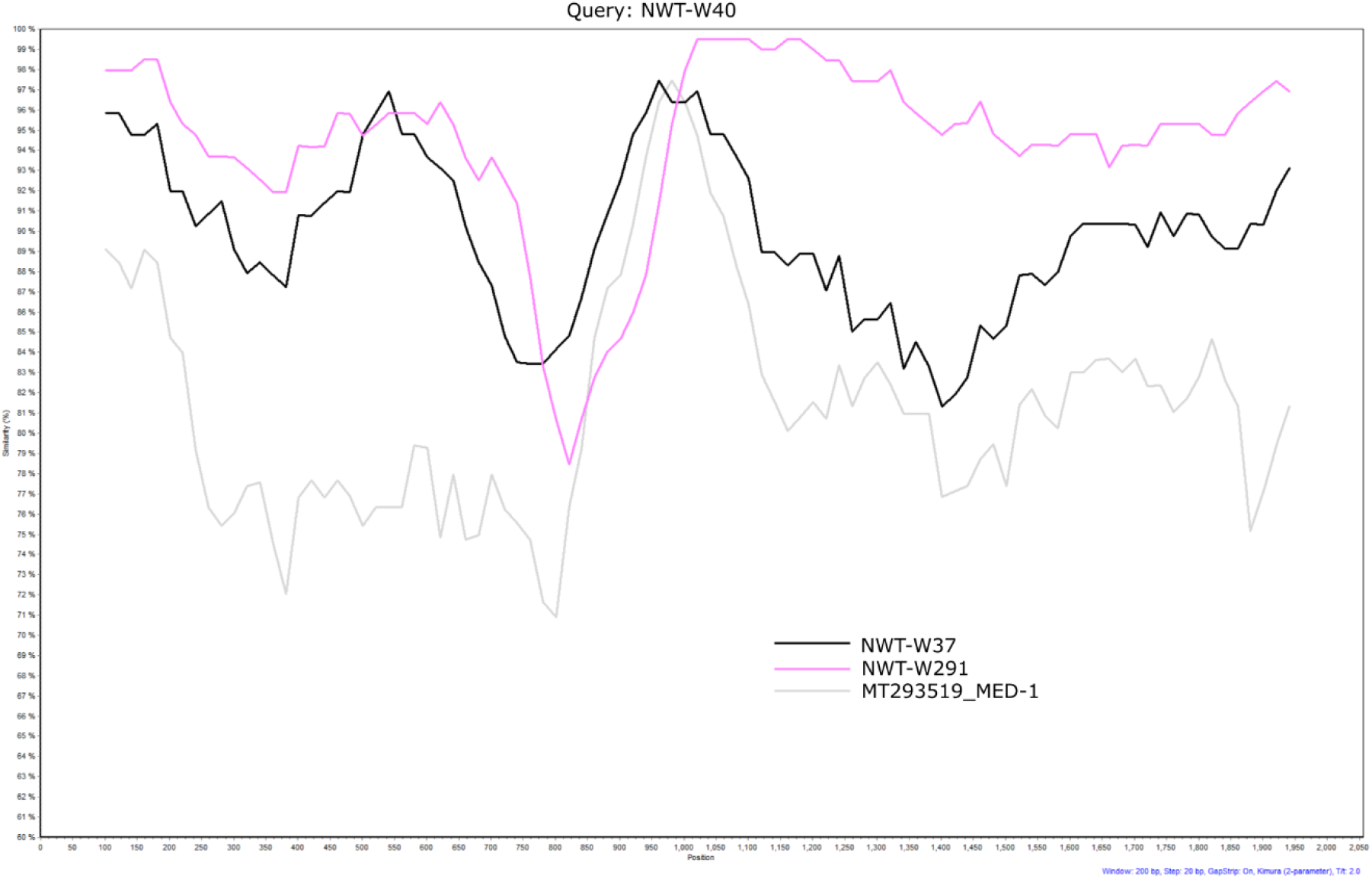
Similarity plot showing the recombinant nature of NWTW-40. The graph shows the percentage identity (y-axis) across the whole genome (x-axis) between strain NWTW-40 (query) and fox circoviruses NWTW-37 and NWTW-291 and canine circovirus MED-1.

## Acknowledgments

The authors would like to acknowledge logistical support from GNWT ECC personnel (Kandis Villebrun, Madison Hurst, Stefan Goodman, Robert Mulders, Kelsey Stewart, and Karl Cox) and local harvesters. MC would also like to acknowledge Dr. Marta Colaneri for insightful conversations about statistical methods.

Preprint version 2 of this article has been peer-reviewed and recommended by Peer Community In Infections (https://doi.org/10.24072/pci.infections.100200; Jean-Francois Guégan, 2024).

## Funding

This work received funding from the Ocean Frontier Institute (OFI) and Canada First Research Excellence Fund (CFREF) through the Visiting Fellowship Program, awarded to MC.

## Conflict of interest disclosure

The authors declare that they comply with the PCI rule of having no financial conflicts of interest in relation to the content of the article. One author (MC) is a recommender for PCI Microbiology and PCI Infections.

## Author contribution

Conceptualization: MC; Data curation: MC; Formal analysis: MC; Funding acquisition: MC, LEL, SCD, ASL; Investigation: MC, AVLK; Methodology: MC; Project administration: MC; Resources: HDC, HF; Visualization: MC; Writing – original draft: MC; Writing – review and editing: AVLK, GF, HDC, LEL, HF, SCD, ASL.

## Data, scripts, code, and supplementary information availability

Sequences obtained in this study are available in GenBank under accession numbers PP438803-PP438871. Metadata are available online at https://doi.org/10.5281/zenodo.10775336; Canuti et al., 2024.

## Notes

https://doi.org/10.5281/zenodo.10775336

### Competing Interest Statement

The authors have declared no competing interest.

### Summary of Updates

The PCIInfections badge certifying that the article has been peer-reviewed and recommended by PCIInfection with the recommendation link has been added.

